# An *in vivo* examination of Dynein-Cargo complex formation

**DOI:** 10.64898/2026.08.02.742322

**Authors:** Phylicia Allen, Hannah Neiswender, Wen Lu, Margot Lakonishok, Rajalakshmi Veeranan-Karmegam, Jessica Pride, Vladimir I. Gelfand, Graydon B. Gonsalvez

## Abstract

Long-range intracellular transport relies on microtubule motors. This process is particularly important in large cells such as neurons and oocytes. While transport towards the plus-end of microtubules utilizes many kinesins, minus-end transport is largely mediated by a single motor, cytoplasmic dynein. Activation of dynein requires the large dynactin complex as well as a cargo adaptor. How dynein, dynactin, and adaptors assemble *in vivo*, particularly within specialized tissues such as the *Drosophila* egg chamber remains unclear. In the current study, we defined the dynein interactome in *Drosophila* egg chambers using *in vivo* proximity biotin ligation. Our findings suggest that Bicaudal-D (BicD) is the principal adaptor responsible for activating dynein and linking it with cargo in this tissue. We also identified Centrocortin (Cen) as a dynein adaptor in egg chambers. However, unlike BicD, loss of Cen did not affect dynein localization or apparent activation. To more specifically analyze adaptor-dependent assembly and cargo transport, we examined dynein light intermediate chain (Dlic) mutants known to impair adaptor binding. As expected, these mutants disrupted the BicD-dynein interaction. However, Cen remained associated with the dynein/dynactin complex in the mutant background, suggesting that Cen engages the motor by a different mechanism. Finally, live imaging of microtubules revealed that even when adaptor binding is compromised, dynein-driven microtubule gliding can still deliver nurse cell-derived cargo into the oocyte, albeit with reduced efficiency. Collectively, our results reveal multiple, mechanistically distinct routes for adaptor association with dynein *in vivo* and indicate that redundant processes can sustain cargo transport during oogenesis.

## INTRODUCTION

Long distance cargo transport within eukaryotic cells is predominantly mediated by microtubule motors. Transport towards the plus-end of microtubules involves one of several kinesins (Hirokawa et al., 2009; Yildiz, 2025). Mammals encode 45 different kinesins and the majority of these are plus-end directed motors (Hirokawa et al., 2009; Yildiz, 2025). By contrast, a single motor, cytoplasmic dynein-1 (hereafter dynein), is responsible for the majority of minus-end directed cargo transport (Canty and Yildiz, 2020; Redwine et al., 2017). Although all cells utilize microtubule motors to transport cargo, this process is especially important in large cells such as neurons. Consequently, mutations in microtubule motors or their accessory factors result in a variety of neurodegenerative and neurodevelopmental diseases (Berth and Lloyd, 2023; Franker and Hoogenraad, 2013). Another type of large cell that relies on microtubule motors for efficient cargo transport is the oocyte (Bastock and St Johnston, 2008; Goldman and Gonsalvez, 2017). For instance, the polarity of the *Drosophila* oocyte and future embryo relies on the microtubule motor mediated transport of *oskar*, *bicoid* and *gurken* mRNAs (Berleth et al., 1988; Ephrussi et al., 1991; Kim-Ha et al., 1991; Neuman-Silberberg and Schupbach, 1993).

The dynein motor exists as a dimer and is composed of several subunits: dynein heavy chain, dynein intermediate chains, dynein light intermediate chains, and three types of dynein light chains (Canty and Yildiz, 2020; Redwine et al., 2017). In the absence of bound cargo, dynein is present in an auto-inhibited conformation known as the phi particle (Torisawa et al., 2014; Zhang et al., 2017). Dynein activation requires dynactin, which is itself a large multi-subunit complex, consisting of several accessory proteins assembled around an actin-like filament (Gill et al., 1991; Schroer and Sheetz, 1991). Association of the dynactin complex with the motor induces conformational changes than are thought to partially activate dynein (Chowdhury et al., 2015; Zhang et al., 2017). Initial studies suggested that the stability of this complex, as well as its processive movement along microtubules, required incorporation of an activating cargo adaptor (McKenney et al., 2014; Schlager et al., 2014). All known activating cargo adaptors contain a coiled-coil domain of variable length, and structural studies have shown that this domain is incorporated between the dynactin complex and the dynein motor (McKenney et al., 2014; Olenick and Holzbaur, 2019; Schlager et al., 2014) (Fig. 1A). These initial studies were conducted using truncated activating adaptors that lacked cargo and were often assembled in the absence of microtubules (McKenney et al., 2014; Schlager et al., 2014). Under these conditions, formation of the dynein-dynactin-adaptor complex was inefficient and required a substantial stoichiometric excess of adaptor protein. By contrast, Rao and colleagues recently demonstrated that inclusion of microtubules greatly enhances formation and stability of the dynein-dynactin complex even in the absence of activating adaptors (Rao et al., 2026). Gillies and colleague also demonstrated that microtubule-bound dynein can independently associate with either dynactin or an activating cargo adaptor (Gillies et al., 2025). Whether these recent findings reflect the mechanism by which the dynein motor complex assembles *in vivo* remains an open question.

**Figure 1:**
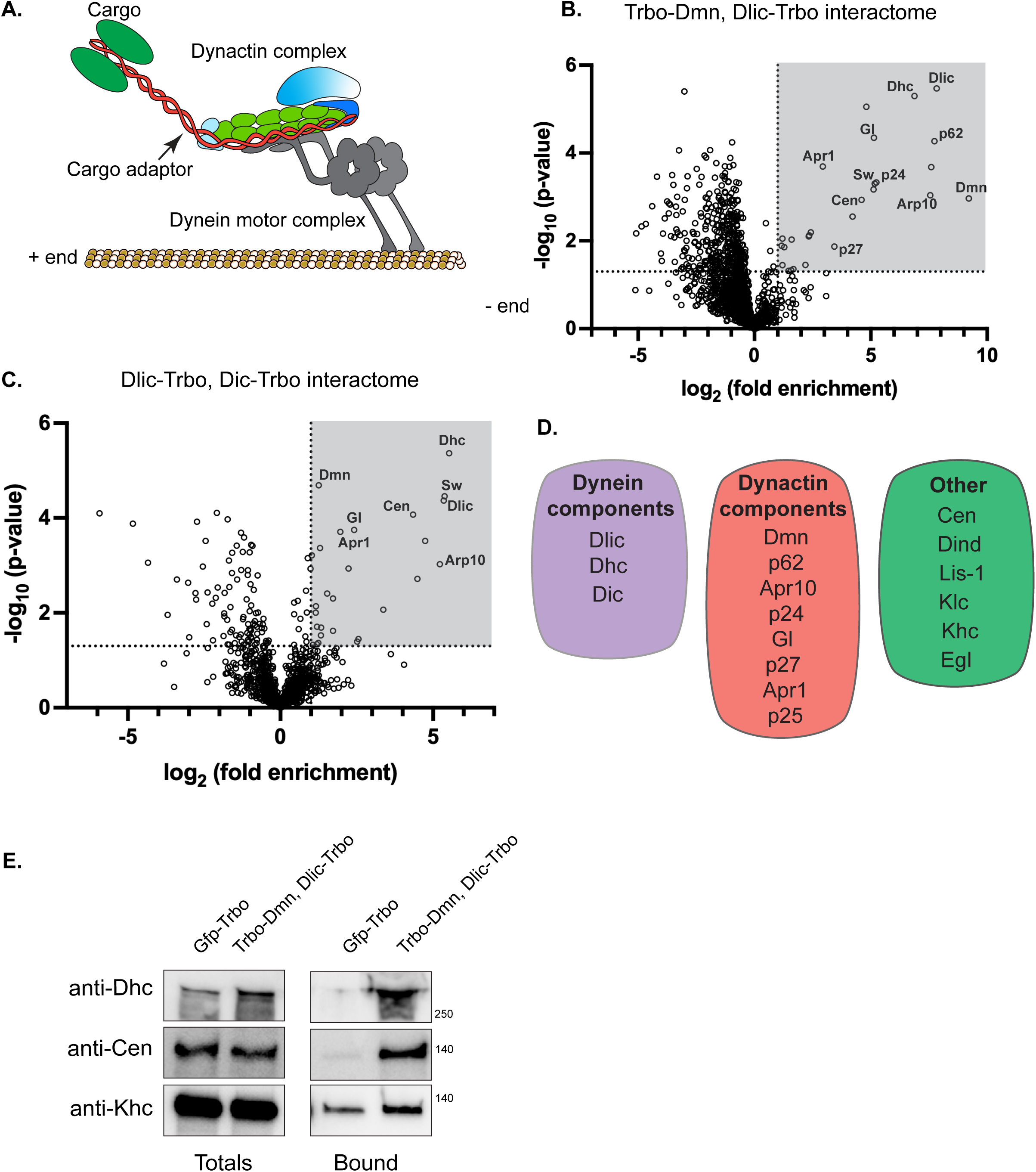
Defining the Dynein interactome. (A) Schematic of the dynein motor. (B) Volcano plot of the Trbo-Dmn/Dlic-Trbo interactome versus GFP-Trbo as a control. The shaded grey box represents proteins that were enriched at least two-fold with the dynein complex and have a p value of at least 0.05. (C) Volcano plot of the Dic- Trbo/Dlic-Trbo interactome versus GFP-Trbo as a control. The specific interacting proteins are shown in the shaded grey box. (D) Select candidates identified in the dynein-dynactin interactomes. The purple box indicates recovered dynein component, the orange box represents recovered dynactin components, and the green box indicates additional candidates that have a functional connection with the dynein/dynactin complex. (E) Validation of top hit in the Trbo- Dmn/Dlic-Trbo interactome. GFP-Trbo served as a negative control. Biotinylated proteins were purified using streptavidin beads and analyzed by western blotting using antibodies against Dhc, Khc, or Cen.

An excellent model to address this topic is the *Drosophila* egg chamber. Oogenesis in *Drosophila* initiates when a single germline stem cell undergoes four rounds of mitotic divisions with incomplete cytokinesis. This results in the formation of a sixteen-cell cyst. One of these cells differentiates as the oocyte, whereas the rest adopt a nurse cell fate (Bastock and St Johnston, 2008; Spradling, 1993). Because the oocyte is halted in meiosis, it is transcriptionally quiescent. As such, growth and maturation of the oocyte depend on transport of factors from the nurse cells into the oocyte via structures known as ring canals (Bastock and St Johnston, 2008). During early and mid-stages of oogenesis, microtubule minus-ends are enriched within the oocyte (Nashchekin et al., 2021; Theurkauf, 1994; Theurkauf et al., 1993; Theurkauf et al., 1992). Thus, transport of mRNAs, proteins, vesicles, and organelles from the nurse cells to the oocyte depend on dynein activity. Loss of dynein activity early in oogenesis results in failure to specify an oocyte, whereas disruption of dynein activity at later stages results in the mis-localization of critical polarity determining mRNAs and proteins (Clark et al., 2007; Goldman and Gonsalvez, 2017; McGrail and Hays, 1997; Mische et al., 2007).

Bicaudal-D (BicD) is the primary adaptor responsible for linking dynein with mRNA cargo in the egg chamber. BicD links these transcripts with dynein via the RNA binding protein Egalitarian (Egl) (Dienstbier et al., 2009; Goldman et al., 2019; Lu et al., 2022; Mach and Lehmann, 1997). In the absence of cargo, BicD is also present in an auto-inhibited conformation due to intramolecular interactions between its N and C termini (Hoogenraad and Akhmanova, 2016; Olenick and Holzbaur, 2019). Egl bound to mRNA cargo interacts with the C-terminal cargo binding domain of BicD, disrupting the intramolecular interaction and enabling BicD to engage the dynein motor (Goldman et al., 2019; McClintock et al., 2018; Sladewski et al., 2018). At present, it is unknown whether cargo adaptors other than BicD also bind and activate dynein in the egg chamber. Furthermore, it is also unknown whether defective adaptor binding results in destabilization of the dynein-dynactin complex and a loss of cargo transport into the oocyte.

To answer these questions, we used proximity biotin ligation to define the dynein interactome in the fly egg chamber. Our findings indicate that in addition to BicD, a recently identified adaptor known as Centrocortin (Cen) is also associated with the dynein complex (Zein-Sabatto et al., 2024). However, unlike BicD, loss of Cen appears to have no effect on the localization and activation of dynein. Furthermore, because dynein light intermediate chain (Dlic) has been shown to bind all known adaptors, we analyzed specific mutations in Dlic known to compromise adaptor binding (Celestino et al., 2019; Lee et al., 2018). As expected, mutant Dlic was defective for interaction with BicD. By contrast, these mutations did not affect the association of Cen with the dynein motor, suggesting that Cen incorporates into the motor by a different mechanism. Lastly, we demonstrate that even under conditions of reduced adaptor binding, dynein mediated microtubule gliding was able to deliver cargo into the oocyte, albeit at a reduced rate.

## RESULTS

### Defining the Dynein interactome

This study aimed to identify the activating adaptors associated with dynein in *Drosophila* egg chambers and to elucidate the mechanisms underlying motor activation for transport. Defining the dynein interactome using traditional purification strategies is challenging because motor-adaptor interactions are weak and transient. To overcome this limitation, we employed *in vivo* proximity biotin ligation using the promiscuous biotin ligase TurboID (Branon et al., 2018). We have used this approach successfully in previous studies to identify the interactome of Egalitarian, the protein that links RNA cargo with the dynein motor via BicD (Baker et al., 2021).

Published studies in mammalian HEK cells demonstrated that Dynein light intermediate chain (Dlic) and Dynein intermediate chain (Dic) fused to a biotin ligase can identify activating adaptors (Redwine et al., 2017). Therefore, we fused the *Drosophila* homologs of these genes with TurboID at the C-terminus (Dlic-Trbo and Dic-Trbo). We also generated flies expressing the p50/Dmn component of the dynactin complex fused to TurboID at the N-terminus (Trbo-Dmn). The positioning of TurboID was based on the known three-dimensional structure of the motor complex and was selected to optimize detection of dynein-interacting proteins (Chowdhury et al., 2015; Redwine et al., 2017; Urnavicius et al., 2015). These constructs were expressed under the control of the alpha-tubulin 67C promoter, which is expressed exclusively in the germline. The constructs also contained a FLAG tag to enable visualization of their *in vivo* localization. All three constructs were appropriately expressed in the germline and enriched within the oocyte, indicating proper dynein-mediated transport (Supplemental Fig. 1A-C). Consistent with the localization pattern of the fusion proteins, biotinylated proteins were also enriched within the oocyte (Supplemental Fig. 1A-C).

We used flies expressing GFP-TurboID (GFP-Trbo) under the same maternal promoter as our control. To maximize labeling of the dynein-dynactin complex and increase the likelihood of identifying potentially novel interacting partners, including activating adaptors, the proteomics experiments were performed using flies co-expressing a combination of either Dlic-Trbo and Trbo- Dmn or Dlic-Trbo and Dic-Trbo. Biotinylated proteins were purified using streptavidin-conjugated magnetic beads and identified using mass spectrometry. The entire experiment was conducted using three independent biological replicates. Proteins enriched at least two-fold in the dynein/dynactin samples with a p-value ≤0.05 were considered dynein-associated or proximal proteins (Fig. 1B, C).

This approach efficiently recovered our bait proteins, Dlic and Dic, as well as dynein heavy chain (Dhc) (Fig. 1D, purple box; Supplemental Tables 1 and 2). Most components of the dynactin complex were also recovered (Fig. 1D, orange box; Supplemental Tables 1 and 2). Notably, we identified the heavy and light chains of the kinesin-1 motor (Khc and Klc) in our dynein/dynactin interactome (Fig. 1D, green box; Supplemental Tables 1 and 2). This finding was not unexpected, as both motors are thought to function together in oocyte polarization (Brendza et al., 2002; Duncan and Warrior, 2002; Januschke et al., 2002) and may therefore be present in the same transport complex. Additionally, we recovered Lis1, which is involved in dynein activation (Canty and Yildiz, 2020; Htet et al., 2020; Reck-Peterson et al., 2018), and Diamond (Dind), a protein involved in dynactin complex formation and stability (Zhang et al., 2024) (Fig. 1D, green box; Supplemental Tables 1 and 2). Another prominent component in the dynein/dynactin interactome was the recently identified adaptor, Centrocortin (Cen) (Zein-Sabatto et al., 2024). The mammalian homologs of Cen, CDR2 and CDR2L, were shown to bind dynein and functions in organization of endoplasmic reticulum (ER) sheets (Teixeira et al., 2025). To validate these results, we repeated the experiment using either GFP-Trbo flies or flies co-expressing Trbo-Dmn and Dlic-Trbo. Biotinylated proteins were purified and analyzed using available antibodies against Dhc, Khc, and Cen. Consistent with our proteomics results, all three proteins were specifically enriched in the dynein/dynactin sample (Fig. 1E).

Numerous studies have shown that dynein function in the female germline relies on BicD (Goldman et al., 2019; Lu et al., 2022; Mach and Lehmann, 1997). Thus, we were surprised that our dynein/dynactin interactome did not contain a statistically significant enrichment of BicD. However, we recovered Egl (Fig. 1D, green box; Supplemental Tables 1 and 2), which links mRNA cargo with dynein via BicD (Dienstbier et al., 2009). The presence of Egl in the dynein/dynactin interactome suggests that our inability to detect BicD is most likely due to the three-dimensional conformation of the complex and the limited labeling surface on BicD that is exposed to TurboID within the context of the dynein motor.

### Characterization of Cen

Most studies involving Cen have examined the function of this protein in *Drosophila* embryos (Kao and Megraw, 2009; Ryder et al., 2020). Given that we recovered Cen as a component of the dynein/dynactin interactome in egg chambers, we examined its localization and function in this tissue. GFP-Cen was expressed in the female germline using a nanos-Gal4 driver. In early and mid-stage egg chambers, GFP-Cen was highly enriched within the oocyte, wherein the protein localized to puncta adjacent to the oocyte cortex (Fig. 2A, B). In stage10 egg chambers, the puncta were localized along the anterior cortex (Fig. 2C), in a pattern reminiscent of the localization of *bicoid* mRNA, another cargo of the dynein motor (Berleth et al., 1988). Thus, the localization of GFP-Cen is consistent with dynein-mediated transport.

**Figure 2:**
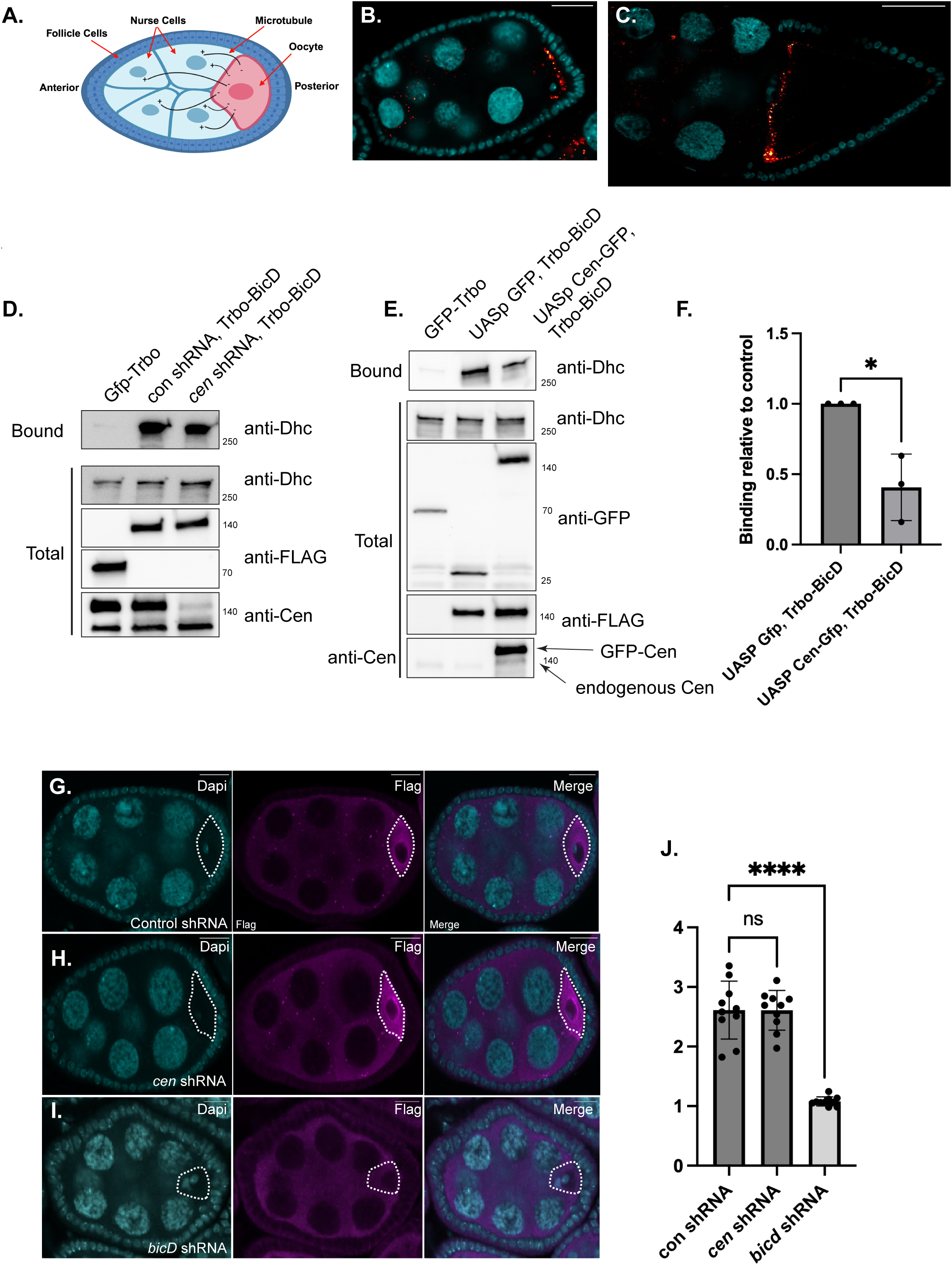
Characterization of Cen. (A) Schematic of a stage 6 *Drosophila* egg chamber. (B-C) Ovaries were dissected from flies expressing Cen-GFP using the nanos-Gal4 driver. The egg chamber were counterstained with DAPI to visualize nuclei. The signal for Cen is show using a red to white lookup table (LUT). A stage 6 egg chamber is shown in panel B, and a stage 10 egg chamber is shown in panel C. Cen in highly enriched in the oocyte in stage 6 chambers and along the anterior cortex of the oocyte in stage 10 egg chambers. (D) Ovarian lysates were prepared from flies expressing Trbo-BicD together with either control shRNA or *cen* shRNA. A strong maternal tubulin Gal4 driver was used to deplete Cen. GFP-Trbo served as a negative control. Biotinylated proteins were purified using streptavidin beads and analyzed by western blotting using the indicated antibodies. The amount of Dhc biotinylated and precipitated by Trbo-BicD was unchanged upon depletion of Cen. (E) Ovarian lysates were prepared from flies expressing Trbo-BicD together with either GFP alone or Cen-GFP. GFP and Cen-GFP were over-expressed using a strong maternal tubulin Gal4 driver. Biotinylated proteins were purified and analyzed by western blotting using the indicated antibodies. The amount of Dhc biotinylated by Trbo-BicD was reduced in strains expressing GFP- Cen. (F) The binding experiment in E was quantified from three independent biological replicates. An unpaired t test was used for this analysis. * p ≤ 0.05. (G-I) Egg chambers expressing a control shRNA (G), *cen* shRNA (H) or *bicD* shRNA (I) in a Dlic_WT background were processed for immunofluorescence and probed with anti-FLAG antibody (magenta) to assess dynein enrichment in the oocyte. The egg chambers were counterstained using DAPI to visualize nuclei (cyan). Depletion of Cen has no effect on Dlic oocyte localization. By contrast, the oocyte enrichment of Dlic is greatly reduced upon depletion of BicD. The oocyte is indicated by dashed lines. (J) Quantification of Dlic oocyte enrichment. A one-way Anova was used for this analysis by comparing the oocyte enrichment of Dlic in the *cen* and *bicD* depletion strains to the control. **** p ≤ 0.0001.

Similar to BicD, Cen also contains a CC1 box motif that is thought to be critical for engaging the dynein motor (Zein-Sabatto et al., 2024). We therefore wondered whether the two adaptors compete for binding dynein. A strain expressing BicD tagged with N-terminal TurboID (Trbo-BicD) was used to test this. These strains expressed either a control shRNA or an shRNA targeting *cen*. Expression of *cen* shRNA effectively depleted the target protein (Fig. 2D, Supplemental Fig. 2A). However, despite the almost complete loss of Cen, the amount of dynein brought down by Trbo- BicD was relatively unchanged (Fig. 2D). If the two adaptors were in competition, we would have expected Trbo-BicD to bring down substantially more dynein upon Cen depletion. We also tested for adaptor competition by examining the interaction between Trbo-BicD and dynein in strains over-expressing GFP-Cen. Over-expression of Cen using a maternal alpha-Tubulin driver had only a modest effect in reducing the amount of dynein brought down by Trbo-BicD (Fig. 2E, F), despite the very high level of GFP-Cen expression (Fig. 2E, compare GFP-Cen to endogenous Cen). Thus, although at very high levels, Cen is able to displace BicD from the dynein motor, the two adaptors do not appear to compete for dynein binding when expressed at endogenous levels. In fact, a small amount of GFP-Cen appeared to be specifically biotinylated by Trbo-BicD, suggesting the possibility that the two adaptors might be present in the same complex (data not shown). To test this more directly, we examined biotinylated pellets from flies expressing either GFP-Trbo or BicD tagged with TurboID on the N and C-terminus (Trbo-BicD and BicD-Trbo). In comparison to the control, a small amount of endogenous Cen was detected in the Trbo-BicD and BicD-Trbo pellets (Supplemental Fig. 2B). Collectively, these results suggest that BicD and Cen do not compete for binding dynein, and that they might even be present in the same complex.

We next determined whether Cen was required for activating dynein for transport into the oocyte. As expected, Dlic with a FLAG tag was enriched within the oocyte in strains expressing a control shRNA (Fig. 2G). The oocyte enrichment of Dlic-FLAG was essentially unchanged upon depletion of Cen (Fig. 2H, J). By contrast, depletion of BicD resulted in a dramatic reduction of Dlic-FLAG within the oocyte (Fig. 2I, J). This is consistent with published results implicating BicD as the primary adaptor for activating dynein in the female germline (Lu et al., 2022). Thus, although Cen appears to be associated with the dynein/dynactin complex *in vivo*, its loss does not appear to significantly compromise dynein activation. This is consistent with the finding that *cen* null alleles are viable and fertile whereas loss of dynein results in lethality (Gepner et al., 1996; Kao and Megraw, 2009).

### Dlic mutants and adaptor binding

A conserved helix within the C-terminus of Dlic has been shown to contact all known mammalian activating adaptors (Celestino et al., 2019; Lee et al., 2018). Mutation of conserved residues within this stretch of amino acids to alanine compromises adaptor binding and cargo transport (Fig. 3A, red arrows) (Celestino et al., 2019; Lee et al., 2018). In *c. elegans*, mutation of these conserved residues phenocopies *dlic* nulls (Celestino et al., 2019). We therefore wished to analyze the activity of these mutants in egg chambers. A wild-type Dlic transgene and three mutant alleles were generated, Dlic_FF, Dlic_LL and Dlic_LFFLL. In each case, the corresponding amino acids were mutated to alanine. Consistent with published results, shRNA-mediated depletion of Dlic results in an oogenesis block (Fig. 3B) (Lu et al., 2022). This phenotype could be rescued by expressing wild-type Dlic in the shRNA background (Fig. 3C, C’). Because the shRNA sequence targets the 3’UTR of endogenous Dlic, expression of the transgenic construct, which contains a distinct 3’UTR, is not affected by the shRNA. Surprisingly, the Dlic mutants also rescued the oogenesis defect (Fig. 3D-F). There was a delay in growth of the oocyte in strains expressing the Dlic mutants (Fig. 3C’, D’, E’, F’ and G). However, by stage10, the size of the oocyte was similar between all strains (Fig. 3H). In addition to a transient defect in oocyte size, there were also subtle defects in the localization of the oocyte marker, Orb (Lantz et al., 1994). In strains expressing wild-type Dlic, Orb was highly enriched within the oocyte (Fig. 3C). Orb was also oocyte enriched in strains expressing the Dlic mutants. However, the level of Orb signal in the nurse cells was higher in the mutants compared to strains expressing wild-type Dlic. In addition, Orb frequently accumulated in puncta within the nurse cell cytoplasm in strains expressing the Dlic mutants (Fig. 3D’, E’ and F’, arrows).

**Figure 3:**
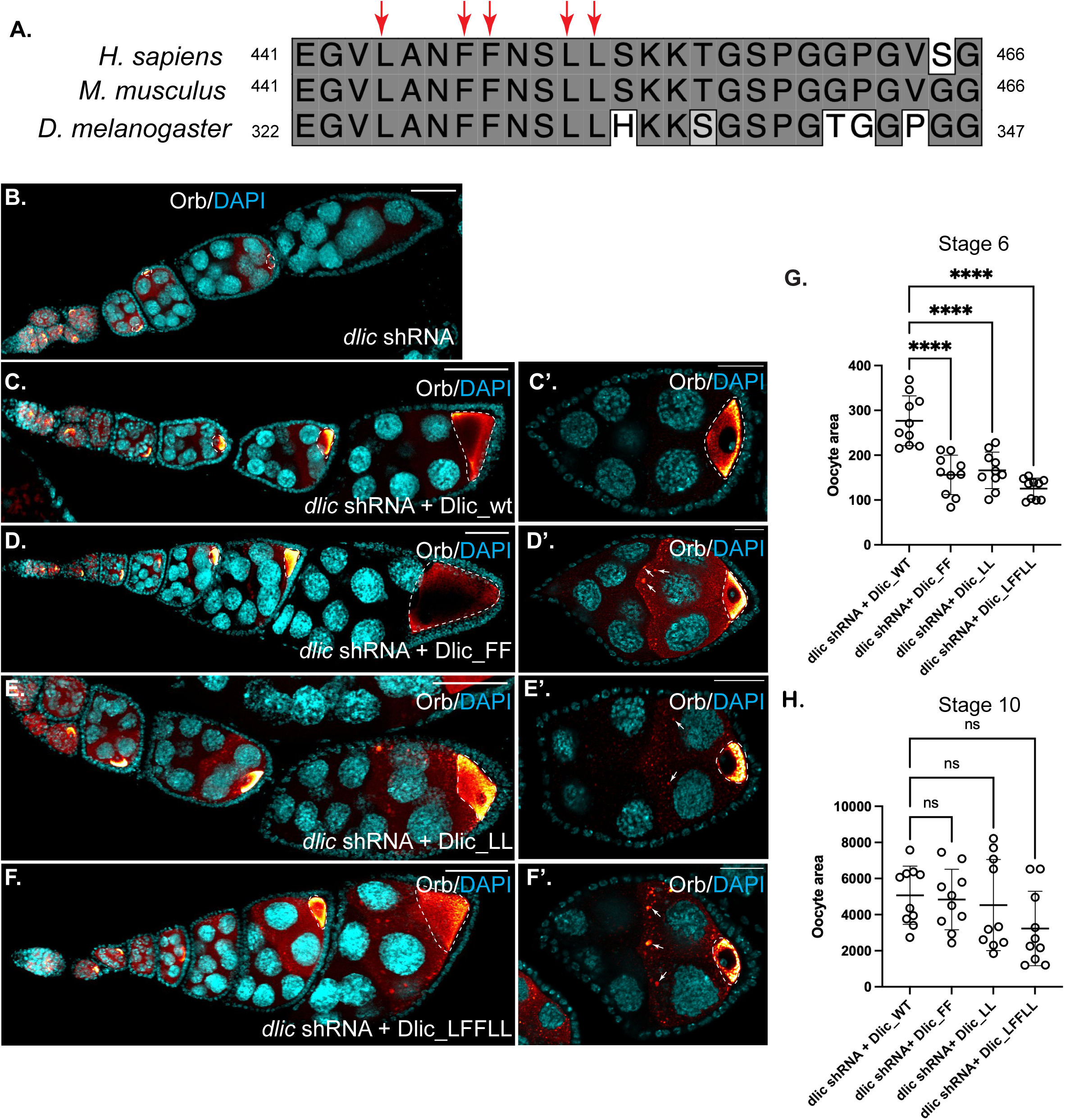
Analysis of adaptor binding defective Dlic mutants. (A)Sequence alignment of conserved residues within helix 1 of Dlic across species: *H. sapiens*, *M. musculus*, and *D. melanogaster*. The critical residues that have been shown to be important for adaptor binding are indicated with red arrows. (B) Ovaries were dissected from flies expressing an shRNA against *dlic*. The egg chambers were processed for immunofluorescence using an antibody against Orb (red to white LUT) and were counterstained with DAPI (cyan). Depletion of Dlic results in an oogenesis arrest. (C-F). Ovaries were dissected from flies co-expressing the *dlic* shRNA as well as either Dlic_WT (C), Dlic_FF (D), Dlic_LL (E) or Dlic_LFFLL mutants. For each mutant the indicated residue was changed to alanine. Dlic_WT as well as the mutants were able to rescue the oogenesis defect caused by depletion of endogenous Dlic and Orb was correctly localized within the oocyte. Puncta of Orb could also be detected in the nurse cell cytoplasm of strains expressing the Dlic mutants (arrows). At early stages there was a delay in growth of the oocyte in the Dlic mutant backgrounds (C’, D’, E’, and F’). By stage 10, the size of the oocyte was similar between strains expressing wild-type and mutant Dlic. (G) Quantification of oocyte size in stage 6 egg chambers. (H) Quantification of oocytes size in stage 10 egg chambers. A one-way Anova was used for this analysis where the size of the oocyte in strains expressing wild-type Dlic was compared to the mutants. **** p ≤ 0.0001, ns = not significant.

Based on published findings showing that mammalian Dlic mutants were defective for binding activating adaptors (Celestino et al., 2019; Lee et al., 2018), we were surprised to find that similar mutations in fly Dlic were able to rescue the oogenesis defect. We therefore examined whether the mutant alleles of Dlic were able to bind activating adaptors in the fly germline. For this experiment, wild-type or mutant FLAG-Dlic was expressed in a background in which endogenous Dlic was depleted. These strains also expressed Trbo-BicD. As expected, Dhc and wild-type Dlic were efficiently biotinylated and precipitated by Trbo-BicD (Fig. 4A). By contrast, the interaction between Trbo-BicD and dynein was compromised in all three Dlic mutant backgrounds (Fig. 4A- C). Similar results were obtained using strains expressing another dynein adaptor, Hook, also tagged with TurboID (Trbo-Hook) (Supplemental Fig. 2C). Our findings are therefore consistent with published results using mammalian Dlic and demonstrate that mutation of critical residues within fly Dlic disrupt binding to activating adaptors.

**Figure 4:**
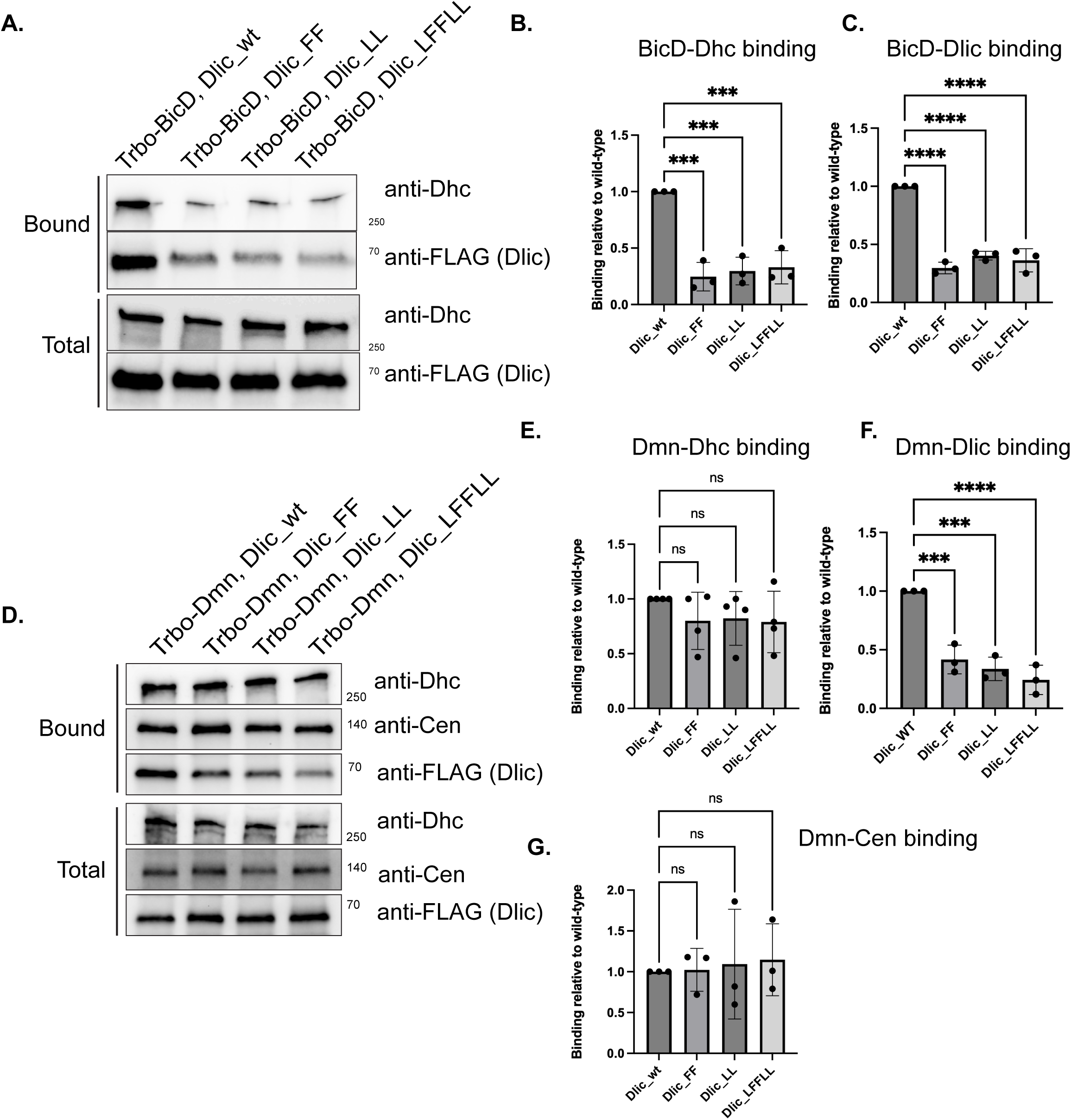
Dynein-cargo complex formation in Dlic wild-type and mutant background. (A) Ovarian lysates were prepared from flies co-expressing Trbo-BicD and *dlic* shRNA. The flies also expressed either wild-type Dlic or the indicated mutants. Biotinylated proteins were purified and analyzed by western blotting and probed with either Dhc or FLAG antibodies. (B-C) The binding experiment between Trbo-BicD and Dhc (B) or Dlic (C) was repeated in triplicate and quantified. The biotinylation of Dhc and Dlic are reduced in strains expressing the Dlic mutants. A one-way Anova was used for this analysis where the binding of Trbo-BicD with Dhc and Dlic was compared between strains expressing wild-type Dlic and the mutants. ***p ≤ 0.001, ****p ≤ 0.0001. (D) Ovarian lysates were prepared from flies co-expressing Trbo-Dmn and *dlic* shRNA. The flies also expressed either wild-type Dlic or the indicated mutants. Biotinylated proteins were purified and analyzed by western blotting using the indicated antibodies. (E-G) Quantification of the binding experiment shown in panel D. A one-way Anova was used for this analysis where the binding of Trbo-Dmn with Dhc (E), Dlic (F) and Cen (G) was compared between strains expressing wild-type Dlic and the mutants. ns = not significant, ***p ≤ 0.00, ****p ≤ 0.001.

Early *in vitro* studies suggested that the stability of the dynein-dynactin complex requires the association of activating adaptors (McKenney et al., 2014; Schlager et al., 2014). By contrast, more recent studies that examined complex assembly in the presence of microtubules, indicate that the dynein-dynactin complex is stable even in the absence of bound adaptors (Rao et al., 2026). The availability of the Dlic mutants, which disrupts adaptor binding *in vivo*, gave us an opportunity to distinguish between these scenarios. Strains that expressed Trbo-Dmn along with either wild-type or mutant Dlic, and in which endogenous Dlic was depleted, were used for this experiment. The amount of Dhc that was biotinylated and precipitated by Trbo-Dmn was unaffected in the mutant Dlic background (Fig. 4D, E). This suggests that the dynein-dynactin interaction remains relatively stable even in the presence of mutant Dlic.

Unlike what was observed for BicD and Hook, the amount of Cen that was biotinylated and precipitated by Trbo-Dmn was similar between strains expressing wild-type and mutant Dlic (Fig. 4D, G). Thus, incorporation of Cen into the dynein motor complex appears less sensitive to mutant Dlic in comparison to other activating adaptors. Lastly, even though the mutations in Dlic are within a C-terminal helix that is not involved in interaction with Dhc (Schroeder et al., 2014), the mutants were biotinylated and precipitated less efficiently by Trbo-Dmn in comparison to the wild-type protein (Fig. 4D, F). This suggests that either the mutant alleles adopt a conformation that disrupt their association with dynein, or that adaptor binding by Dlic stabilizes the incorporation of this subunit into the dynein motor.

### Dynein-mediated nurse cell to oocyte transport

To correlate the above biochemical phenotypes with function, we examined the transport of dynein and its cargos into the oocyte. As expected, wild-type Dlic was highly enriched in the oocyte (Fig. 5A, C). By contrast, the oocyte enrichment of Dlic mutants was greatly reduced (Fig. 5B, C, Supplemental Fig. 3A, B). This is consistent with their inefficient incorporation into the dynein motor. Dhc and BicD were also highly enriched in the oocyte in strains expressing wild-type Dlic (Fig. 5D, F, G, I). The oocyte enrichment of both proteins was reduced in strains expressing mutant Dlic, with the phenotype for BicD being less severe than Dhc (Fig. 5E, F, H, I, Supplemental Fig. 3C-F). We next examined two cargoes, ER vesicles and Me31b, both of which were shown to localize to the oocyte in a BicD-dynein dependent manner (Baker et al., 2021). Somewhat surprisingly, the oocyte enrichment of both ER vesicles and Me31b was either unaffected or only modestly affected in the mutant background (Fig. 5J-L, Supplemental Fig. 3G- L). Interestingly however, ER vesicles and Me31b accumulated in large aggregates within the nurse cell cytoplasm in strains expressing the Dlic mutants. In fact, both proteins co-localized within these aggregates (Fig. 5M-O). We observed a similar co-localization between Me31b and Orb in the nurse cell aggregates (Supplemental Fig. 3M-O). One possibility is that these aggregates represent stalled transport particles containing cargos that are transported in a dynein dependent manner. Despite the presence of these aggregates, however, our results suggest that cargo transport into the oocyte remains at least partially active in the Dlic mutant background.

**Figure 5:**
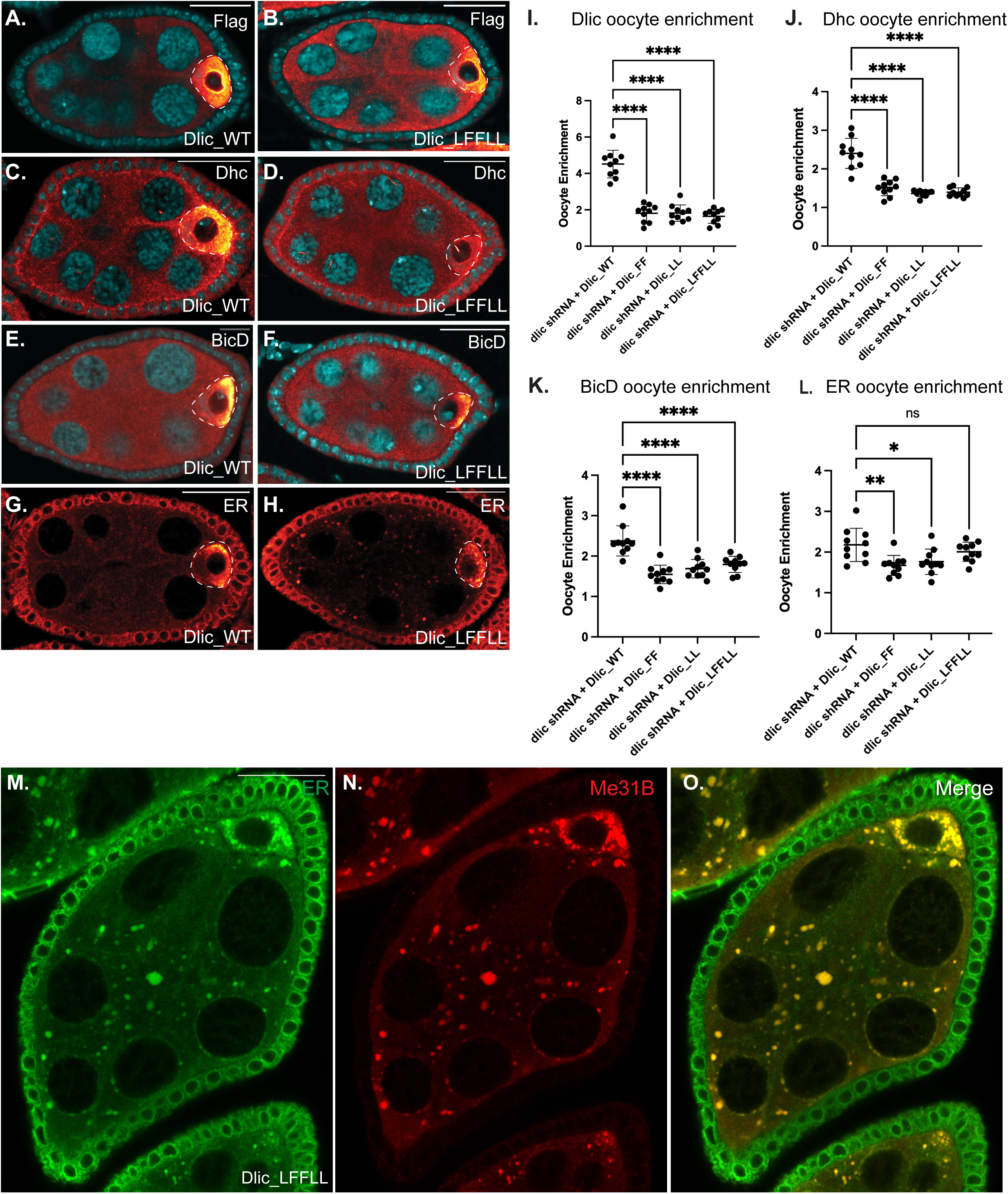
Motor and cargo localization in Dlic wild-type and mutant background. (A, H) Ovaries were dissected from flies co-expressing the *dlic* shRNA and either Dlic_WT or the Dlic_LFFLL mutant. The egg chambers were processed for immunofluorescence using antibodies against FLAG (A, B), Dhc (C, D), BicD (E, F), or an ER marker antibody (G, H). The fluorescent signal for these proteins is shown using the red to white LUT. The egg chambers shown in panels A-F were also counterstained with DAPI (cyan). (I-L) The oocyte enrichment of Dlic (I), Dhc (J), BicD (K) and the ER marker (L) was quantified. A one-way Anova was used for this analysis where the oocyte enrichment of each protein in strains expressing wild-type Dlic was compared to strains expressing the Dlic mutants. ns = not significant, *p ≤ 0.05, **p ≤ 0.01, ****p ≤ 0.001. The oocyte enrichment of Dlic, Dhc and BicD is reduced in strains expressing the Dlic mutants. By contrast, the oocyte enrichment of the ER was either unaffected or only modestly affected in strains expressing the Dlic mutants. (M-O) Ovaries were dissected from flies co-expressing the *dlic* shRNA and the Dlic_LFFLL mutant. The egg chambers were processed for immunofluorescence using antibodies against the ER marker (green) and Me31b. A merged image is also shown. Me31b and ER vesicles co-localize in large nurse cell aggregates. The scale bar is 20 microns.

### Dynein-mediated microtubule gliding remains partially active in the Dlic mutant background

In addition to direct cargo transport, cortically anchored dynein has also been shown to glide microtubules within the nurse cell cytoplasm and from the nurse cells into the oocyte (Lu et al., 2022). This process results in the generation of an advection force that non-specifically moves cargo into the oocyte (Lu et al., 2022). We reasoned that microtubule gliding could function as a redundant mechanism to move cargo such as ER vesicles and Me31b into the oocyte in the Dlic mutant background. We examined this process using a strain in which microtubules were labeled with the EMTB microtubule binding domain fused to GFP. As expected, robust microtubule gliding was observed in strains expressing wild-type Dlic (Fig. 6A, A’, video 1). Microtubule gliding was also observed in strains expressing the Dlic_LFFLL mutant (Fig. 6B, B’, video 2). However, the number of motile microtubules was reduced in the mutant background (Fig. 6C). Furthermore, the velocity of these microtubules as well as their net displacement was also reduced in comparison to the wild-type (Fig. 6D, E). Thus, although microtubule gliding still occurs in the Dlic mutant background, the efficiency of this process is reduced.

**Figure 6:**
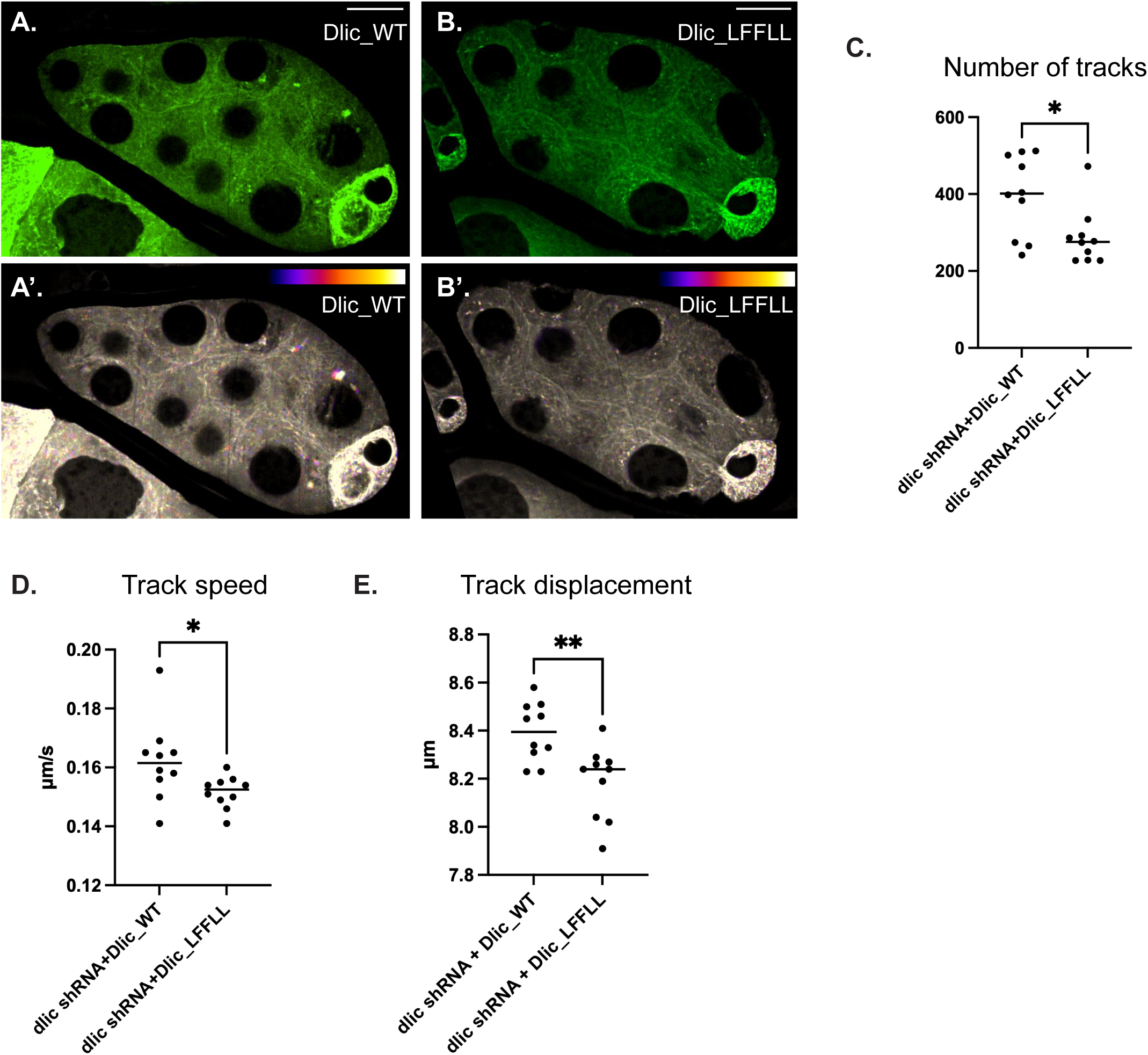
Microtubule gliding remains partially active in the Dlic_LFFLL mutant. (A, B) Ovaries were dissected from flies co-expressing EMTB-3XGFP and the *dlic* shRNA to deplete endogenous Dlic. The strains also expressed either transgenic wild-type Dlic (A) or the Dlic_LFFLL mutant (B). The egg chambers were imaged live to examine microtubule gliding. A temporal color-coded image of the first fifteen frames of the movie is shown in panels A’ and B’. (C-E) The number of motile tracks was quantified between strains expressing wild-type versus the Dlic_LFFLL mutant (C). In addition, the track speed (D) and net track displacement (E) was also quantified between these strains. An unpaired t-test was used for this analysis. *p ≤ 0.05, **p ≤ 0.01. The scale bar is 20 microns. Although microtubule gliding still occurs in the Dlic mutant, the number of motile tracks, the track speed and the track displacement are all reduced.

Collectively, our results suggest that the Dlic-adaptor interaction is critical for formation of a fully active dynein motor complex. However, redundant interactions between the adaptors and the dynein/dynactin complex enable partial dynein-cargo assembly and transport. In addition, microtubule gliding, which also remains partially active in the Dlic mutant background, is able to non-specifically move cargo into the oocyte. Thus, although both processes, direct cargo transport and dynein-mediated gliding, are reduced in the Dlic mutants, the residual activity of both processes is sufficient to deliver cargos into the growing oocyte.

## DISCUSSION

Cargo transport towards the minus-end of microtubules occurs mostly via dynein (Reck-Peterson et al., 2018). Given this reliance on a single motor, cargo adaptors that link dynein with its diverse cellular cargo play an essential role (Olenick and Holzbaur, 2019). Importantly, cargo adaptors also perform another critical function; they convert dynein from an inactive, weakly processive motor, to one that is highly processive (McKenney et al., 2014; Schlager et al., 2014). In addition to cargo adaptors, the large multi-subunit dynactin complex is also required for dynein activation (Gill et al., 1991; Schroer and Sheetz, 1991).

Initial *in vitro* studies suggested that cargo adaptors, which are sandwiched in between dynactin and dynein, stabilize the trimeric complex. However, these studies were performed in the absence of microtubules and cargo, and used truncated adaptors present in large stoichiometric excess (McKenney et al., 2014; Schlager et al., 2014). Recent *in vitro* studies, in which the complexes were assembled under more physiological conditions and in the presence of microtubules, suggest that the dynein/dynactin complex is stable even without cargo adaptors (Rao et al., 2026). Moreover, these studies also demonstrated that adaptors compete for binding to the dynein motor even after a dynein/dynactin/adaptor complex has already been assembled (Rao et al., 2026). Whether these *in vitro* results also hold true for dynein activation and cargo binding *in vivo*, however, remains unknown. Our goal, therefore, was to use a well-characterized *in vivo* model, the *Drosophila* egg chamber, to address this question.

In this study, we employed *in vivo* proximity biotin ligation using TurboID to reveal the composition of the dynein-associated proteome in *Drosophila* egg chambers. Our strategy of fusing TurboID to multiple dynein and dynactin components (Dlic, Dic, and p50/Dmn) proved effective in capturing the dynein interactome. For instance, we recovered most components of the dynactin complex as well as Diamond (also known as Vezatin), a factor required for assembly and stability of this large multi-subunit complex (Zhang et al., 2024). In addition, we also recovered Lis1, a well-known regulator of dynein activity (Markus et al., 2020). Current models suggests that the main function of Lis1 is to modulate the conformation of the auto-inhibited dynein motor such that it more efficiently associates with dynactin and a cargo adaptor, in essence priming the motor for activation (Geohring et al., 2026; Rao et al., 2026). Also present in our dynein interactome was the heavy and light chains of the plus-end motor, kinesin-1. This finding is consistent with earlier studies suggesting that both motors function in a coordinated manner to polarize the fly oocyte (Brendza et al., 2002; Duncan and Warrior, 2002; Januschke et al., 2002), and with findings in mammalian cells that several cargos are transported in a bi-directional manner by associating with opposite polarity motors (Canty et al., 2023; Kendrick et al., 2019; Splinter et al., 2010).

Numerous studies over the past two decades have established a biochemical and functional relationship between BicD and the dynein motor in the fly egg chamber. BicD and dynein are both required for oocyte specification and maintenance, and in later stage egg chambers, BicD links dynein to critical polarity determining mRNAs (Bullock and Ish-Horowicz, 2001; Clark et al., 2007; Dienstbier et al., 2009; Dienstbier and Li, 2009; Goldman et al., 2021; Goldman et al., 2019; Huynh and St Johnston, 2000; Lu et al., 2022; Mach and Lehmann, 1997; McClintock et al., 2018; Neiswender et al., 2021; Sladewski et al., 2018; Vazquez-Pianzola et al., 2017). One objective of this project was to determine whether additional cargo adaptors also associate with dynein in the egg chamber. Surprisingly, the only other cargo adaptor we identified in the dynein interactome was Cen. Cen as well as its mammalian homologs, CDR2 and CDR2L, were recently shown to interact with dynein via an N-terminal CC1 box (Teixeira et al., 2025; Zein-Sabatto et al., 2024), similar to the dynein-interaction motif found within BicD. CDR2 and its paralog CDR2L were required for maintaining proper organization of ER sheets in mammalian cancer cells (Teixeira et al., 2025). The main function attributed for *Drosophila* Cen is in mediating the localization of its own mRNA to the centrosome of embryos and to function along with another protein, Centrosomin, in formation of the embryonic cleavage furrow (Kao and Megraw, 2009; Ryder et al., 2020; Zein-Sabatto et al., 2024).

Based on our identification of Cen in the dynein interactome, and the fact that both BicD and Cen contain a similar CC1 box for dynein interaction, we wondered whether the two adaptors compete for biding to dynein. Although at very high expression level, Cen was able to displace BicD from dynein, the adaptors do not appear to compete for binding dynein when expressed at endogenous levels. In fact, our results suggest that both adaptors might actually be present within the same complex. Whether or not the adaptors are co-transported with dynein, however, remains to be determined. We also find that whereas BicD is required for activating dynein and promoting its transport into the oocyte, Cen was dispensable for this process. Consistent with this notion, flies that are homozygous null for *cen* are viable and fertile (Kao and Megraw, 2009). Our results do not rule out a role for Cen in dynein activation but rather suggests that at least within the context of the egg chamber, the primary adaptor required for dynein-mediated oocyte transport is likely BicD. *In vitro* studies using CDR2 showed that this protein could activate dynein motility, albeit at a much lower level than another dynein adaptor that was used for comparison, JIP3 (Teixeira et al., 2025). Thus, the exact role of *Drosophila* Cen in dynein mediated transport remains to be fully explored.

To further investigate the molecular requirements for adaptor-dynein interactions, we took a genetic approach by mutating a key interaction interface. A critical attachment point between cargo adaptors and dynein appears to be via a conserved interaction with a helical segment of Dlic (Celestino et al., 2019; Lee et al., 2018; Schroeder et al., 2014). Mammalian Dlic has been shown to interact with most activating adaptors and mutation of conserved leucine and phenylalanine residues within this helix disrupts adaptor binding (Celestino et al., 2019; Lee et al., 2018). Thus, to better understand the mechanism of dynein-cargo assembly, we generated similar mutations in fly Dlic. Loss of Dlic in the germline results in oogenesis arrest (Lu et al., 2022). As expected, expression of wild-type Dlic in this background completely restored oogenesis. Surprisingly, however, expression of the Dlic mutants also restored oogenesis, despite a transient slow-down in oocyte growth. The rescue of oogenesis occurs even though the Dlic mutans displayed a substantially reduced interaction with the adaptors BicD and Hook. Consistent with disrupted adaptor binding, the oocyte enrichment of Dlic, Dhc and BicD were all reduced. Recent ultrastructural studies have found that in addition to the Dlic-adaptor interaction, additional contact points exist between the dynein motor and cargo adaptors (d’Amico et al., 2026). These redundant interactions most likely enable residual dynein-adaptor binding in the Dlic mutant background, thus resulting in rescue of oogenesis.

Another unexpected finding was that cargo transport into the oocyte was relatively unaffected in the mutant Dlic background. This suggests that redundant mechanism exist for moving cargo into the oocyte. One potential mechanism that might account for this finding is dynein-mediated microtubule gliding. This occurs via cortically anchored dynein that glides microtubule instead of traditional cargo (Lu et al., 2022). The movement of these microtubules in the viscous nurse cell cytoplasm generates advection forces that non-specifically moves contents into the oocyte. We thus examined microtubule gliding in the Dlic_LFFLL mutant. This revealed that although microtubule gliding still occurred in the mutant background, the process was less efficient. Thus, the Dlic-adaptor interaction is required for efficient microtubule gliding.

Collectively, our results suggest that the Dlic mutants compromise, but do not eliminate, specific cargo transport as well as microtubule gliding. These redundant mechanisms, occurring at a reduced level, appear to be sufficient to rescue the oogenesis defect as well as cargo transport into the oocyte. Similar to our findings with dynein, redundant mechanisms are also responsible for proper cargo localization within the oocyte. For instance, the localization of the posterior determinant *oskar* mRNA is dependent on direct transport via the kinesin-1 motor (Brendza et al., 2000). However, kinesin-1 is also required for microtubule sliding within late-stage oocytes, a process referred to as cytoplasmic streaming (Lu et al., 2016; Palacios and St Johnston, 2002). Cytoplasmic streaming cooperates with direct transport of *oskar* mRNA to ensure robust localization of this cargo to the posterior of the oocyte (Lu et al., 2018). Thus, *Drosophila* appear to have evolved multiple mechanisms to sustain delivery of critical cargo to the growing oocyte and to maintain its correct localization within this large cell.

In summary, our *in vivo* analysis reveals that BicD and Cen both associate with dynein in egg chambers, but only BicD is essential for dynein activation and oocyte localization. Importantly, mutations in the Dlic tail that disrupt adaptor binding compromise, but do not eliminate, dynein function. This partial loss of function is compensated by redundant transport mechanisms, direct cargo transport and microtubule gliding, highlighting the robust strategies *Drosophila* employs to ensure proper oocyte development. These findings bridge *in vitro* biochemical studies with *in vivo* function and demonstrate that cells have evolved multiple fail-safe mechanisms for critical developmental processes.

## ACKNOWLEDGEMENTS

We are grateful to Dr. Timothy Megraw for generously providing the Cen antibody and to Dr. Akira Nakamura for generously providing the anti-Me31b antibody. We are also grateful to Pritha Bagchi for her assistance with the mass-spectrometry analysis. We also thank the Bloomington stock center and the Developmental Studies Hybridoma bank for providing essential fly strains and antibodies. This work was supported by grants from the National Institutes of Health (R35GM145340 to GBG and 2R35GM131752 to VIG) and from the CCBx research program at the Simons Foundation’s Flatiron Institute (to VIG and WL). This work was also supported in part by the Emory Integrated Proteomics Core (RRID:SCR_023530) and the Augusta University Cell Imaging Core (RRID:SCR_026799).

## MATERIALS AND METHODS

### DNA Constructs

TurboID (Trbo) constructs were cloned into the pAttB vector (Bischof et al., 2007). A fragment containing the promoter and 3’UTR for alpha-Tubulin67c as well as the SV40 poly adenylation sequence was generated by gene synthesis and cloned into this vector using the NEB high fidelity assembly kit. Sequences corresponding to the cDNA for either GFP, Dlic, Dic, Dmn, BicD and Hook, also generated by gene synthesis, were then cloned into the pAttB-maternal promoter vector. Next, the TurboID sequence, codon optimized for expression in *Drosophila*, was generated by gene synthesis and cloned into the above vector. This generated maternally driven GFP-Trbo, Dlic-Trbo, Dic- Trbo, Trbo-Dmn, Trbo-BicD, BicD-Trbo, and Trbo-Hook. The Dlic wild-type and mutant constructs (Dlic_FF, Dlic_LL and Dlic_LFFLL) also containing a 3xFLAG tag were generated by gene synthesis and cloned into the above pattB-maternal promoter vector. Gene synthesized fragments were generated by Genewiz/Azenta. All constructs were verified by whole plasmid sequencing from Plasmidsaurus prior to generation of transgenic flies.

### Fly Stocks and crosses

Fly crosses were maintained at 25^0^C. For examination of ovarian phenotypes, mated female flies were dissected at 3 to 4 days of age. Transgenic strains were generated by BestGene Inc. The following lines were generated.

pattB-maternal promoter-GFP-Trbo (injected into Bloomington strain 24485, Chr3, 68E)

pattB-maternal promoter-Dlic-Trbo (injected into Bloomington strain 24485, Chr3, 68E)

pattB-maternal promoter-Dic-Trbo (injected into Bloomington strain 24482, Chr2, 51C)

pattB-maternal promoter-Trbo-Dmn (injected into Bloomington strain 24485, Chr3, 68E)

pattB-maternal promoter-Trbo-BicD (injected into Bloomington strain 24485, Chr3, 68E)

pattB-maternal promoter-Trbo-Hook (injected into Bloomington strain 24485, Chr3, 68E)

pattB-maternal promoter-Dlic_wt (injected into Bloomington strain 24485, Chr3, 68E)

pattB-maternal promoter-Dlic_FF (injected into Bloomington strain 24485, Chr3, 68E)

pattB-maternal promoter-Dlic_LL (injected into Bloomington strain 24485, Chr3, 68E)

pattB-maternal promoter-Dlic_LFFLL (injected into Bloomington strain 24485, Chr3, 68E)

The following stocks were obtained from the Bloomington stock center.

- pUASp-Cen-EGFP (#53751)
- *eb1 shRNA* (#36680, TRiP.HMS01568, used as a control for knock-down experiments)
- *cen* shRNA (#43139, TRiP.GL01480)
- *bicD* shRNA (#35405, TRiP.GL00325)
- mat αtub-Gal4[V37] (#7063)
- nanos-gal4 (#4442)

To analyze the interaction between BicD and dynein shown in Fig.2D, the Trbo-BicD strain was recombined with the mat atub-Gal4 driver. This strain was then crossed to the respective shRNA strain. To examine complex formation upon over-expression of Cen (Fig. 2E), the Trbo-BicD strain recombined with the mat atub-Gal4 driver was crossed to the indicated GFP tagged strains. To examine the localization of Dlic upon depletion of Cen or BicD (Fig. 2G-I), a strain expressing Dlic_wild-type was first recombined with the mat atub-Gal4 driver. This strain was then crossed with the respective shRNA strain. To examine the rescue phenotype of Dlic wild-type and mutants (Fig. 3), the respective Dlic strains were first recombined with the mat atub-Gal4 driver. These strains were then crossed to the *dlic* shRNA strain which targets the 3’UTR of endogenous *dlic* (Lu et al., 2022). To examine dynein-cargo complex formation (Fig. 4), the Trbo-BicD and Trbo- Dmn strains were first recombined with the *dlic* shRNA strain. These strains were then crossed to the Dlic wild-type and mutant strains that were recombined with the mat atub-Gal4 driver. For the microtubule gliding experiment shown in Fig. 6, a strain expressing *EMTB-3XGFP-sqh 3’UTR* (attP40, a gift from Yu-Chiun Wang lab, RIKEN Center for Biosystems Dynamics Research) (Lu et al., 2022) was brought into the background of the mat atub-Gal4 driver. This strain was then crossed to a strain in which either Dlic_WT or Dlic_LFFLL was recombined with the *dlic* shRNA.

### Antibodies

The following antibodies were used: mouse anti-FLAG (Millipore-Sigma, 1:500 for immunofluorescence, 1:5000 for western), mouse anti-Dhc 2C11-2 (Developmental studies hybridoma bank, 1:50 for immunofluorescence, 1:300 for western), anti-Cen C (courtesy of Dr. Timothy Megraw,1:10,000 for western), rabbit anti-Khc (generated inhouse 1:5000 for western), rabbit anti-GFP (Proteintech, 1:1000 for western), mouse anti-Orb (Developmental studies hybridoma bank, 1:30 for immunofluorescence), mouse anti-BicD 1B11 (Developmental studies hybridoma bank, 1:30 for immunofluorescence), mouse anti-BicD 4C2 (Developmental studies hybridoma bank, 1:30 for immunofluorescence), mouse anti-Cnx99A (Developmental studies hybridoma bank, 1:60 for immunofluorescence), rabbit anti-Me31b (courtesy of A. Nakamura, 1:2,000 for immunofluorescence). The following secondary antibodies were used: goat anti-mouse Alexa 488 and 555 (Life Technologies, 1:400) and goat anti-rabbit Alexa 488 and 555 (Life Technologies, 1:400). Biotinylated proteins were detected in localization experiments using Streptavidin-Alexa647 (Life Technologies, 1:1200).

### Purification and analysis of biotinylated proteins

Proteins biotinylated by TurboID were purified from ovarian lysates as previously described (Baker et al., 2021). In brief, lysates were prepared using RIPA buffer (50 mM Tris-cl [pH 7.5], 150 mM NaCl, 1% NP-40, 1 mM EDTA) containing a Halt Protease inhibitor cocktail (Thermo Fischer Scientific). For small scale experiments, 1mg of ovarian lysates was used and incubated overnight with 30ul of High-Capacity Streptavidin Agarose beads (Thermo Fisher Scientific) at 4^0^C. The next day, samples were washed four times with RIPA buffer, bound proteins were eluted in Laemmli buffer and analyzed by western blotting. All western blot images were acquired on a Bio Rad ChemiDoc MP.

For larger scale proteomic experiments, roughly 100 flies were dissected per genotype. 5mg of ovarian lysate from each genotype was used per binding experiment. Lysates were incubated with 100ul Streptavidin magnetic beads (Thermo Fisher Scientific) at 4^0^C overnight. The next day, samples were extensively washed using 1ml of the following: three times with RIPA buffer, three times with high-salt RIPA buffer (50 mM Tris-Cl [pH 7.5], 1 M NaCl, 1% NP-40, 1 mM EDTA), three times with RIPA buffer, and four times with PBS. The entire experiment was conducted using three independent biological replicates. After the final wash, beads were resuspended in 100ul of PBS and shipped to the Emory Integrated Proteomics Core on dry ice. Bound proteins were eluted by trypsin digestion and analyzed by mass spectrometry.

### Mass spectrometry

A published protocol was followed for on bead digestion of proteins (Soucek et al., 2016). Beads were resuspended in digestion buffer (100 mM Tris-HCl), then reduced with 1 mM dithiothreitol (DTT) for 30 minutes at room temperature. Alkylation was subsequently carried out using 5 mM iodoacetamide (IAA) for 30 minutes at room temperature in the dark. Bound proteins were first digested overnight with 1ug of lysyl endopeptidase (Wako) at room temperature, followed by a second overnight digestion with 1ug of trypsin (Promega) at room temperature. The resulting peptides were desalted using an HLB column (Waters) and dried by vacuum.

The data acquisition by LC-MS/MS was adapted from a published procedure (Seyfried et al., 2017). Dried peptides were reconstituted in loading buffer (0.1% trifluoroacetic acid, TFA) and separated on a self-packed 25 cm column (100 µm internal diameter, packed with 1.7 µm Waters CSH beads) coupled to a Dionex 3000 RSLCnano liquid chromatography system. Chromatographic separation was achieved at a flow rate of 1.5 µL/min using the following gradient profile: initial conditions of 1% buffer B (80% acetonitrile, 0.1% formic acid) ramping to 5% over 6 seconds, followed by a 35-minute linear gradient to 23% B, a 6-minute gradient to 35% B, and a final 4-minute wash at 99% B. Mass spectra were acquired on a Q-Exactive HF-X mass spectrometer (ThermoFisher Scientific, San Jose, CA). The instrument was operated in a data-dependent acquisition mode, collecting one full MS scan followed by 20 MS/MS scans per cycle. Full MS scans were performed over a 400–1600 m/z range at a resolution of 120,000 at m/z 200 in profile mode, with an AGC target of 1×10⁶ and a maximum ion injection time of 100 ms. HCD MS/MS spectra were collected at a resolution of 30,000 at m/z 200, using an isolation width of 1.6 m/z, a normalized collision energy of 28%, an AGC target of 2×10⁴, and a maximum ion injection time of 50 ms. Dynamic exclusion was applied to previously sequenced precursor ions for 20 seconds within a 10 ppm window. Precursor ions with unassigned, +1, +6 to +8, or greater than +8 charge states were excluded from fragmentation.

Label-free quantification analysis was adapted from a published procedure (Seyfried, Dammer et al. 2017). Spectra were searched using the search engine Andromeda, integrated into MaxQuant (v. 2.4.2.0), against 2020 Drosophila melanogaster (Fruit fly) Uniprot database (42,676 target sequences). Methionine oxidation (+15.9949 Da), asparagine and glutamine deamidation (+0.9840 Da) and protein N-terminal acetylation (+42.0106 Da) were variable modifications (up to five allowed per peptide); cysteine was assigned as fixed carbamidomethyl modification (+57.0215 Da). Only fully tryptic peptides with up to two miscleavages were considered in the database search. A precursor mass tolerance of ±20 ppm was applied before mass accuracy calibration and ±4.5 ppm after internal MaxQuant calibration. Other search settings included a maximum peptide mass of 6,000 Da, a minimum peptide length of six residues and 0.05-Da tolerance for high resolution MS/MS scans. The FDR for peptide spectral matches, proteins and site decoy fraction was set to 1%. Quantification settings were as follows: match full MS1 peaks between runs; use a 0.7-min retention time match window after an alignment function was found with a 20-min retention time search space. The LFQ algorithm in MaxQuant was used for protein quantitation. The quantitation method considered only razor and unique peptides for protein level quantitation.

This work was supported by the Emory University Emory Integrated Proteomics Core Facility (RRID:SCR_023530).

### Immunofluorescence

Dissected ovaries were fixed in 4% formaldehyde for 20 minutes at room temperature. Egg chambers were then blocked for 1 hour at room temperature using 5% normal goat serum (Thermo Fischer Scientific). Samples were incubated in blocking solution with primary antibody overnight at 4^0^C. The following day, samples were washed twice with PBST (PBS + 0.1% Triton X-100) and incubated with secondary antibody in blocking solution overnight at 4^0^C. The next day, samples were washed 3 times with PBST and then stained with DAPI. The samples were then mounted on slides using Prolong Diamond (Life technologies).

### Microscopy

Imaging experiments described in all figures expect Fig. 6 were performed at the Augusta University Cell Imaging Core (RRID:SCR_026799). Fixed images were captured on an inverted Nikon AXR confocal microscope equipped with the NSPARC detector. For live imaging of microtubule movement in *Drosophila* egg chambers, young mated adult females expressing EMTB-3xGFP and either Dlic_WT or Dlic_LFFLL and depleted of endogenous Dlic were fed dry active yeast for 16–18 hours and then dissected in Halocarbon oil 700 (Sigma-Aldrich, Cat# H8898), as previously described previously (Lu et al., 2023; Lu et al., 2022). Freshly dissected samples were imaged using a Nikon W1 spinning disk confocal microscope equipped with a Yokogawa CSU spinning disk unit with a 50 µm pinhole, a Hamamatsu ORCA-Fusion digital CMOS camera, and a 40× 1.25 N.A. silicone oil-immersion objective. Image acquisition was controlled using Nikon Elements software, and images were collected for 5min at 5sec intervals. Images were processed for presentation using Fiji, Adobe Photoshop, and Adobe Illustrator.

### Quantifications

To calculate the enrichment of Dhc, Dlic, BicD, ER vesicles, and Me31b, the mean pixel intensity of each protein in the oocyte was divided by the mean pixel intensity of the same protein in the nurse cells in individual egg chambers. This analysis was performed using Fiji. To quantify microtubule gliding, the surface feature in Imaris10.2 (Oxford Instruments) was used along with machine learning in order define the microtubule structure. For particle tracking, the autoregressive motion algorithm was used with a max distance of 3 microns and max gap size of 3. Tracks that had a net displacement length of less than 6 microns were filtered out to remove particles that displayed mostly diffusive motion. Graphpad Prism10 was used to perform statistical analysis and for graph creation.

**Supplemental figure 1:** (A-C) Ovaries were dissected from strains expressing Dlic-Trbo (A), Dic- Trbo (B) or Trbo-Dmn (C). The ovaries were processed for immunofluorescence using a FLAG antibody (cyan). The ovaries were also incubated with Alexa674 conjugated streptavidin to reveal the localization of biotinylated proteins (magenta). A merged image is also shown. The scale bar is 20 microns. All three fusion constructs are enriched within the oocyte (dashed lines).

**Supplemental figure 2:** (A) Ovaries were dissected from flies expressing a control shRNA or an shRNA targeting Cen. Lysates were prepared and analyzed by western blotting using the indicated antibodies. (B) Ovaries were dissected from flies expressing either GFP-Trbo, Trbo- BicD or BicD-Trbo. Biotinylated proteins were purified and analyzed by western blotting using an antibody against Cen. (C) Ovaries were dissected from flies expressing Trbo-Hook and either wild-type Dlic or the indicated mutants. Biotinylated proteins were purified and analyzed by western blotting using the FLAG antibody.

**Supplemental figure 3:** (A-H) Ovaries were dissected from flies expressing *dlic* shRNA to deplete endogenous Dlic as well as transgenic Dlic_FF or Dlic_LL. The egg chambers were processed for immunofluorescence using antibodies against FLAG (A, B), BicD (C, D), Dhc (E, F), or the ER marker antibody (G, H). The fluorescent signal for these proteins is shown using a red to white LUT. (I-L) Ovaries were dissected from flies expressing *dlic* shRNA as well as transgenic Dlic_WT (I), Dlic_FF (J), Dlic_LL (K) or Dlic_LFFLL (L). The egg chambers were processed for immunofluorescence using an antibody against Me31b. The fluorescent signal proteins is shown using a red to white LUT. (M-O) Ovaries were dissected from flies expressing *dlic* shRNA and Dlic_LFFLL and the egg chambers were processed for immunofluorescence using antibodies against Orb (green) and Me31b (red). A merged image is also shown. The scale bar is 20 microns.

**Supplemental table 1:** Excel file listing the candidates identified in the Trbo-Dmn/Dlic-Trbo interactome.

**Supplemental table 2:** Excel file listing the candidates identified in the Dlic-Trbo/Dic-Trbo interactome.

**Video 1:** Live imaging of flies expressing EMTB-GFP in the wild-type Dlic background.

**Video 2:** Live imaging of flies expressing EMTB-GFP in the Dlic_LFFLL mutant background.

