## Supplementary figures and images for "An *in vivo* examination of Dynein-Cargo complex formation"

### Supplemental figure1

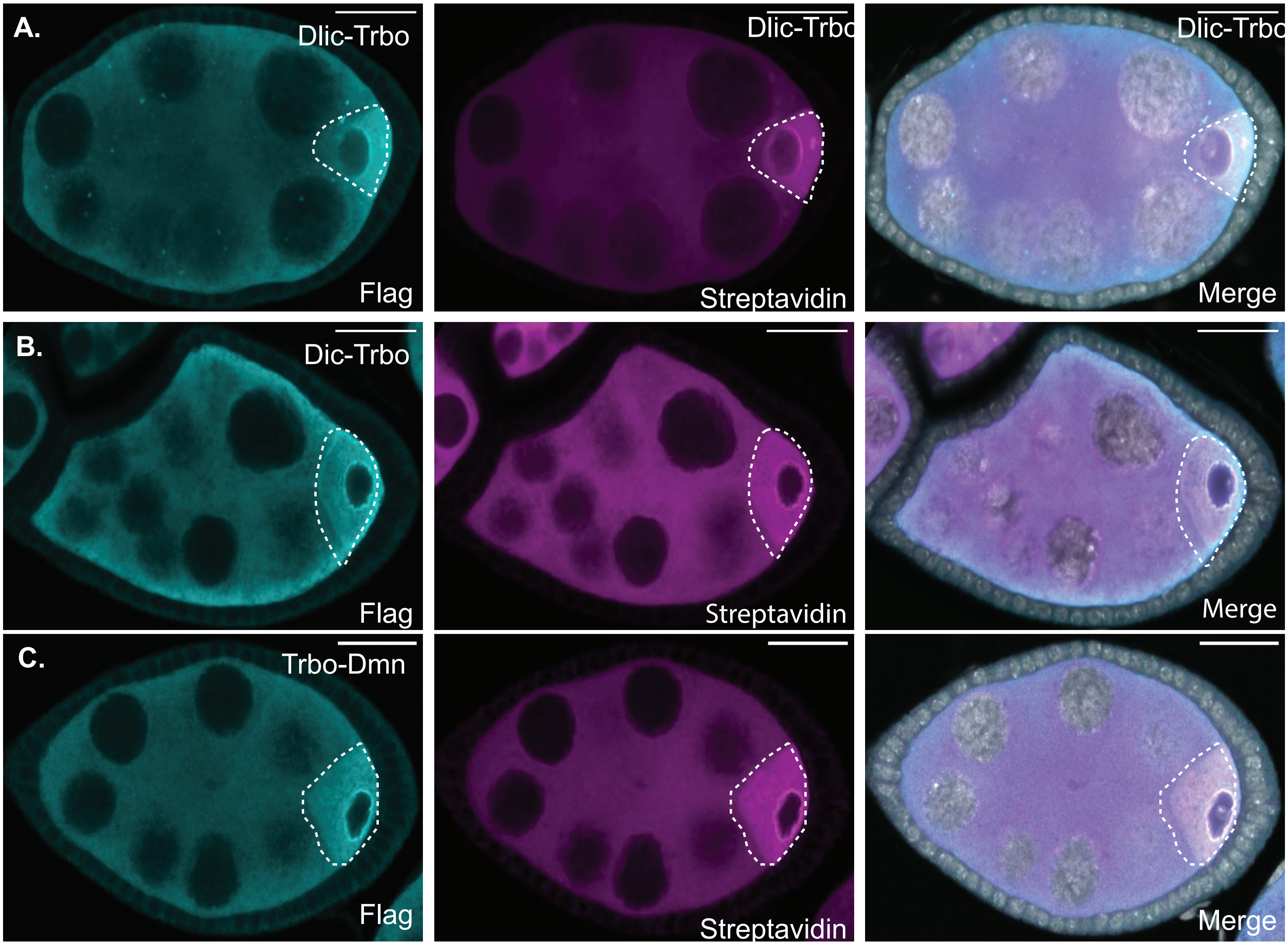


Supplemental Figure 1

### Supplemental figure2

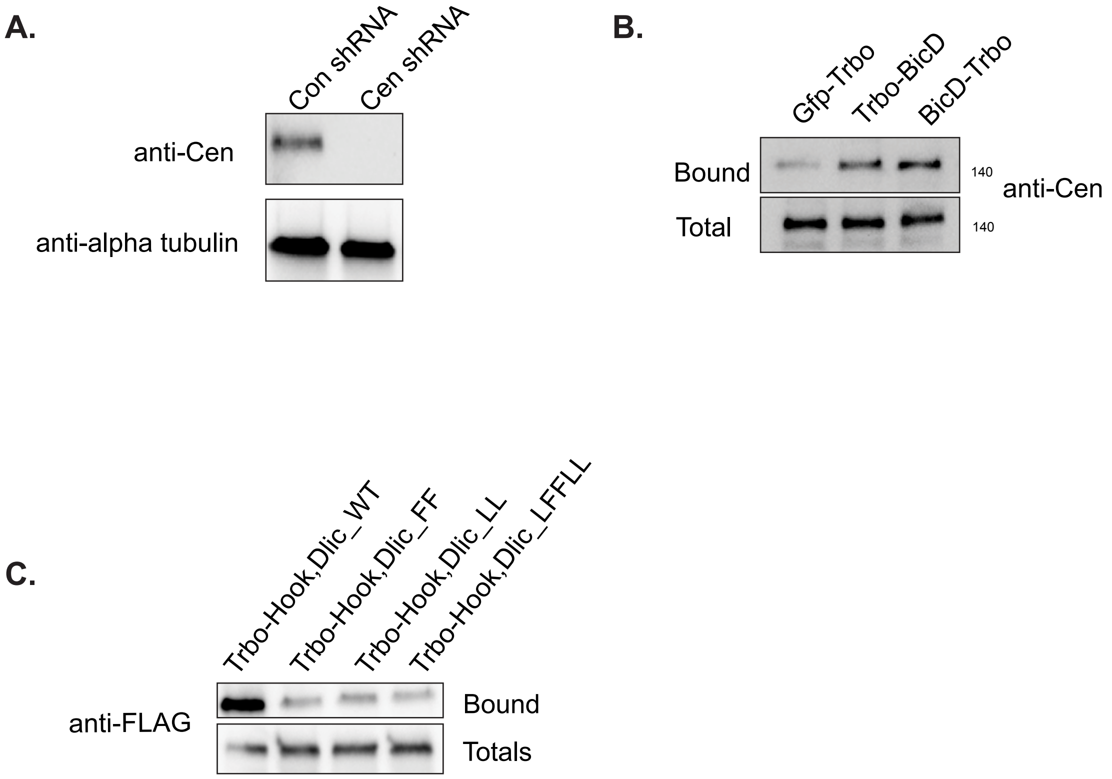


Supplemental Figure 2

### Supplemental figure3

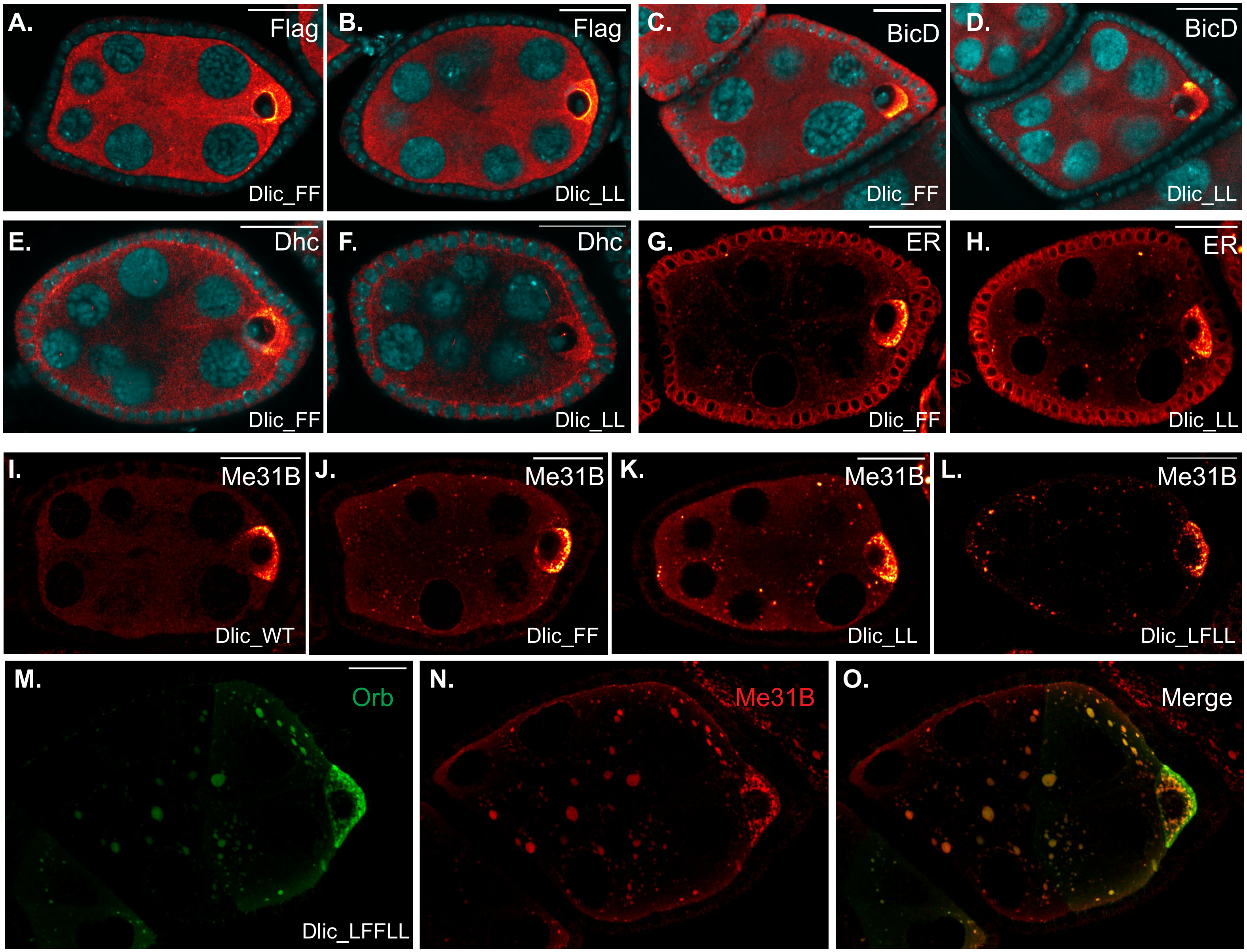


Supplemental Figure 3
